# Deep learning-based detection of murine congenital heart defects from µCT scans

**DOI:** 10.1101/2024.04.06.588383

**Authors:** Hoa Nguyen, Audrey Desgrange, Amaia Ochandorena-Saa, Vanessa Benhamo, Sigolène M. Meilhac, Christophe Zimmer

**Affiliations:** Institut Pasteur, Université Paris Cité, Imaging and Modeling Unit, F-75015 Paris, France; Université Paris Cité, Imagine - Institut Pasteur Unit of Heart Morphogenesis, INSERM UMR1163, F-75015, Paris, France; Rudolf Virchow Center, University of Würzburg, Germany

**Keywords:** Congenital heart defects, micro-computed tomography, mouse, 3D-CNN, segmentation, diagnosis, deep learning

## Abstract

Congenital heart defects (CHD) result in high morbidity and mortality rates, but their origins are poorly understood. Mouse models of heart morphogenesis are required to study the pathological mechanisms of heart development compared to normal. In mouse fetuses, CHD can be observed and detected in 3D images obtained by thoracic micro-computed tomography (μCT). However, diagnosis of CHD from μCT scans is a time-consuming process that requires the experience of senior experts. An automated alternative would thus save time, empower less experienced investigators and could broaden analysis to larger numbers of samples.

Here, we describe and validate an approach based on deep learning to automatically segment the heart and screen normal from malformed hearts in mouse μCT scans. In an initial cohort, we collected 139 μCT scans from thorax and abdomen of control and mutant perinatal mice. We trained a self-configurating neural network (nnU-Net) to segment hearts from body μCT scans and validated its performance on expert segmentations, achieving a Dice coefficient of 96%. To identify malformed hearts, we developed and trained a 3D convolutional neural network (CNN) that uses segmented μCT scans as inputs. Despite the relatively small training data size, our diagnosis model achieved a sensitivity, specificity (for a 0.5 threshold), and area under the curve (AUC) of 92%, 96%, and 97% respectively, as determined by 5-fold cross-validation.

As further validation, we analyzed two additional cohorts that were collected after the model was trained: a ‘prospective’ cohort, using the same experimental protocol as the initial cohort, and containing a subset of its genotypes, and a ‘divergent’ cohort in which mice were subjected to a different treatment for heart arrest (cardioplegia) and that contained a new mouse line. Performance on the prospective cohort was excellent, with a sensitivity of 92%, a specificity of 100%, and an AUC of 100%. Performance on the divergent cohort was moderate (sensitivity: 69%, specificity: 80% and AUC: 81%), but was much improved when the model was finetuned on (a subset of) the cohort (sensitivity: 79%, specificity: 88% and AUC: 91%). These results showcase our model’s robustness and adaptability to technical and biological differences in the data, highlighting its usefulness for practical applications.

In order to facilitate the adoption, adaptation and further improvement of these methods, we built a user-friendly Napari plugin (available at napari-hub.org/plugins/mousechd-napari) that allows users without programming skills to utilize the segmentation and diagnosis models and re-train the latter on their own data and resources. The plugin also highlights the cardiac regions used for the diagnosis. Our automatic and retrainable pipeline, which can be employed in high-throughput genetic screening, will accelerate diagnosis of heart anomalies in mice and facilitate studies of the mechanisms of CHD.

## Introduction

Congenital malformations are a leading cause of pregnancy arrest^1^ and perinatal lethality in developed countries^2^, and the most frequent malformations are those of the heart^3^. While genetics is thought to play a major role in cardiac developmental defects, the specific genetic causes are known only for a fifth of the patients^4^, mainly monogenic mutations associated with syndromes or cardiomyopathies. Polygenic inheritance or gene-environment interactions are believed to underlie a large part of the other cases, including structural heart defects. However, owing to the rarity of any given combination of mutations in human pathology, dissecting the specific contribution of the hundreds of genes involved in heart development is a daunting task.

The mouse is a valuable model to study heart morphogenesis thanks to the relative proximity of heart structures and developmental patterns between humans and mice^5^. Genetically-modified mouse strains contribute to exploring genotype-phenotype relationships underlying congenital defects. Data collected from these strains has been used to build up extensive databases such as those assembled by the Mouse-Genome Informatics (MGI)^6^ or the International Mouse Phenotyping Consortium (IMPC)^7^.

Several imaging techniques enable the precise capture and identification of 3D cardiovascular structures. These include micro-computed tomography (µCT), magnetic resonance imaging (MRI), and optical coherence tomography. Thanks to technical refinements and improvement in spatial resolution, mouse embryogenesis is now characterized at earlier stages and higher throughput^8^. However, extracting meaningful insights from the vast amounts of data generated by these techniques has become a new challenge for embryologists, as the examination of individual 3D images is a complex and time-consuming process. For example, manual segmentation of organs or tumors by clinicians requires on the order of ∼10 minutes per scan ^9–11^; detailed segmentation of a mouse heart takes about 30 minutes and phenotyping takes between 20 min and 1 hour per heart and requires extensive anatomical training and knowledge of the medical nomenclature of heart defects.

Therefore, automating the analysis of 3D cardiac imaging data holds the potential to increase the throughput of anatomic characterizations. A reliable automated segmentation of the heart volume would allow extracting detailed quantitative measurements of the heart structure, and an automated screen of normal versus malformed hearts could speed up and potentially improve the identification of organ anomalies, while adequate visualization tools could highlight variations in morphological features characteristic of diseased samples.

Although there are previous reports of automated segmentation and classification of CHD in mice^12–16^, they have several limitations. First, segmentation and analysis algorithms, when publicly available, typically require coding skills to be used, and hence call for user-friendly alternatives^14,15^. Secondly, previous work on analyzing morphological phenotypes of developing mice in µCT data has leveraged non-linear registration of CT-scans towards a population-averaged image, and/or separate analyses of voxel intensities, tensor-based morphometry, and comparison of whole-organ volumes with wild-type mice^12,14,15^. The use of non-linear registration makes morphological analysis potentially susceptible to errors caused by anatomical structures unrelated to the heart. The reliance on handcrafted diagnostic features (i.e. features defined by a human expert) leaves open the possibility that more pertinent, less intuitive features can be determined from the data using machine learning techniques, potentially leading to better diagnosis. Deep learning methods, including convolutional neural networks (CNNs)^17,18^, iteratively refine and optimize features for a given task. Therefore, image features learned by CNNs can emerge as new phenotypic descriptors that have not been explicitly defined. Additionally, by jointly analyzing multiple features, neural networks may enable the diagnosis of subtle phenotypes involving several features whose departure from normality may have been insufficient for reliable diagnosis when considered separately.

In this study, we describe a 3D-CNN-based approach to automatically segment heart volumes from a µCT-scan of the thorax and abdomen of mouse fetuses and classify cardiac anatomy as normal or abnormal. We validate the segmentation and diagnosis tools quantitatively on perinatal mice, at E(embryonic day) 18.5 and at birth. Moreover, we provide these methods as a plugin for the widely popular Napari^19^, allowing users to apply our heart segmentation and classification tools without programming experience. Importantly, this plugin can also be used to finetune the classification model on the user’s own data. The proposed tools should be instrumental for fast and objective screening of mouse samples for the study of heart morphogenesis and congenital cardiac defects.

## Materials and Methods

### Initial cohort and diagnosis

In our initial cohort, we used mouse fetal samples with heterotaxy, a disease which includes defects in all cardiac segments. The cohort combines *Hoxb1^Cre/+^;Nodal^flox/-^* conditional mutants, *Rpgrip1l^-/-^* mutants, which have a fully penetrant phenotype (implying that 100% of mutant hearts are abnormal) ^20^ ^5,21^, as well as littermate wild-type or heterozygous controls, and *Ift20^-/+^;Nodal^-/+^*double heterozygotes with no cardiac phenotype. Samples were generated, genotyped and imaged as described previously^5,20^. Heart defects were diagnosed independently by two experts (an embryologist and a clinician) on μCT and high resolution episcopic microscopy (HREM) images^5^.

The initial cohort included 139 μCT scans of 139 individual mice (Figure 1), comprised of 56 scans of mice at E18.5 and 83 mice at birth (P0). Out of the 139 mice, 38 (27%) (Figure 1b) suffered from congenital heart defects (CHD). These CHDs belonged to 5 major non-exclusive categories: apex malposition (dextrocardia, mesocardia), atrial situs defects (right isomerism, left isomerism or situs inversus of atrial appendages), septal defects (complete atrioventricular septal defects), ventricle malposition (L-LOOP), and malposition of the great arteries (double outlet right ventricle, transposition of the great arteries), as shown in the Venn diagram of Figure 1b. (One scan with CHD was excluded from the Venn diagram only (Figure 1b) because the type of anomaly was difficult to interpret). Among the 37 remaining hearts, 36 hearts displayed at least two malformations. Septal defects were present in all 38 abnormal hearts.

**Figure 1:**
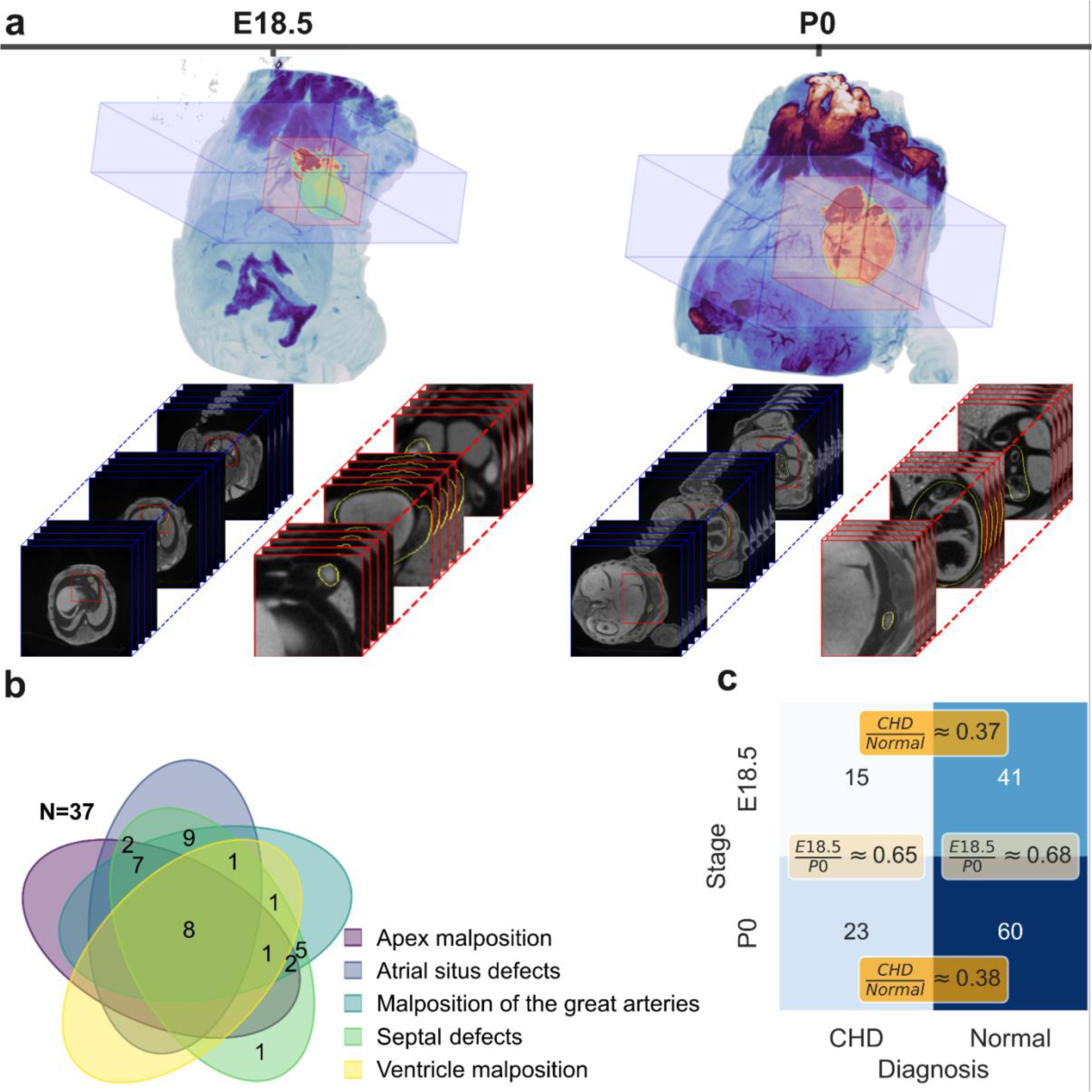
Micro-computed tomography (µCT) images of mouse fetuses and phenotyping of congenital heart defects. **a)** μCT scans of the thorax and abdomen of mouse fetuses at 18.5 days of development (E18.5, left) and at birth (P0, right) displayed as 3D renderings (top) or stacks of 2D slices (bottom). Blue bounding boxes show the set of 2D thoracic slices containing the heart. Red bounding boxes are framed around the heart. The corresponding 2D image stacks are shown with matching color frames below. Yellow contours on each slice outline the segmented heart. **b)** Venn diagram showing the distribution of five types of heart malformations for N=37 mice with heterotaxy. Out of the 37 cases, 36 cases have at least two anomalies, and eight cases exhibit all five types of anomalies simultaneously. **c)** Contingency table for diagnostic label (CHD or normal) and developmental stage (E18.5 or P0).

Hearts at different developmental stages naturally differ in size and morphology (Figure 1a), thus developmental stage is a potential confounder for the classification of heart anomalies. To alleviate this risk, we analyzed scans with equal proportions of CHD and normal hearts at both stages (37% for E18.5 and 38% for P0) (Figure 1c).

### Heart segmentation with nnU-Net

In order to avoid technical confounders, we standardized raw μCT scans to the same axial view (Left – Posterior – Superior: LPS) and to isotropic voxels of size 0.02 mm x 0.02 mm x 0.02 mm. After this preprocessing, we trained a nnU-Net^22^ segmentation model to isolate the heart from the body scans. The nnU-Net is a self-configurating neural network framework for biomedical image segmentation based on the U-Net^23^ architecture, which has demonstrated high performance and robustness for a large number of data types^22^. The nnU-Net pipeline automatically sets up policies for data preprocessing, network architectures, hyperparameters for training and postprocessing, based on data fingerprints. The nnU-Net framework automatically splits training scans into five folds and applies a cross-validation strategy to train five distinct models. The final segmentations are obtained from the ensemble averages of the probabilities predicted by these five models.

For training, the nnU-Net requires a segmentation ground-truth. To define this ground truth, the heart was manually outlined using Imaris (Bitplane) by an embryologist in each 2D slice containing the heart. This was done in a subset of μCT scans consisting of 23 scans with normal hearts (12 mice at E18.5 and 11 at P0) and 17 scans with CHD (10 mice at E18.5 and 7 at P0), totaling 40 scans and 6,261 slices. From these 40 scans, 28 scans served for training the nnU-Net, while 12 scans were kept for testing - Table 1 details the categories and developmental stages for each data set in the initial cohort. This data split guaranteed that all stages and phenotypes were represented in both training and testing data.

**Table 1.** Data split for training and testing nnU-Net segmentation framework (initial cohort). Table entries indicate the number of scans from the initial cohort used for training and testing the nnU-net. CHD: Congenital Heart Defect; E18.5, P0: developmental stages at which scans were taken.

| Stage | Train |  | Test |  |
| --- | --- | --- | --- | --- |
|  | Normal | CHD | Normal | CHD |
| E18.5 | 10 | 9 | 2 | 1 |
| P0 | 7 | 2 | 4 | 5 |
| Total | 28 |  | 12 |  |

Qualitative assessment of the segmentations was performed using Fiji, comparing the outlines of the segmentation with the µCT-scan. Imperfect heart segmentation corresponds either to over-segmentation (i.e. inclusion of non-cardiac regions), or under-segmentation (i.e. exclusion of heart regions).

### Classification model and training procedure

To classify hearts into normal or CHD, we developed a dedicated 3D CNN model that uses the nnU-net based segmentation above to crop out the heart from the fetus body and mask out non-heart pixels (Figure 2a). To help compensate for the low number of training images, we took one out of every five slices along the z-axis to form a 3D image from each μCT scan, then repeated this process by shifting the first slice by one slice four times, thus artificially increasing the number of training images by five-fold (at the expense of a five-fold reduction of axial resolution) (Figure 2b). For training, these images were re-interpolated linearly along the z-axis to obtain volumes with isotropic voxels. For testing, however, the trained model was applied to the entire set of slices contained in the cropped heart region, without subsampling. Due to differences in heart volumes, the resulting 3D images had different sizes (193 ± 25, 154 ± 21, 165 ± 20 voxels). We resized all images using linear interpolation to a common size of 64 x 64 x 32 voxels before feeding them as input to the classification CNN. To further diversify the training data, we applied Gaussian noise and random crops to these standardized 3D images during training on-the-fly.

**Figure 2:**
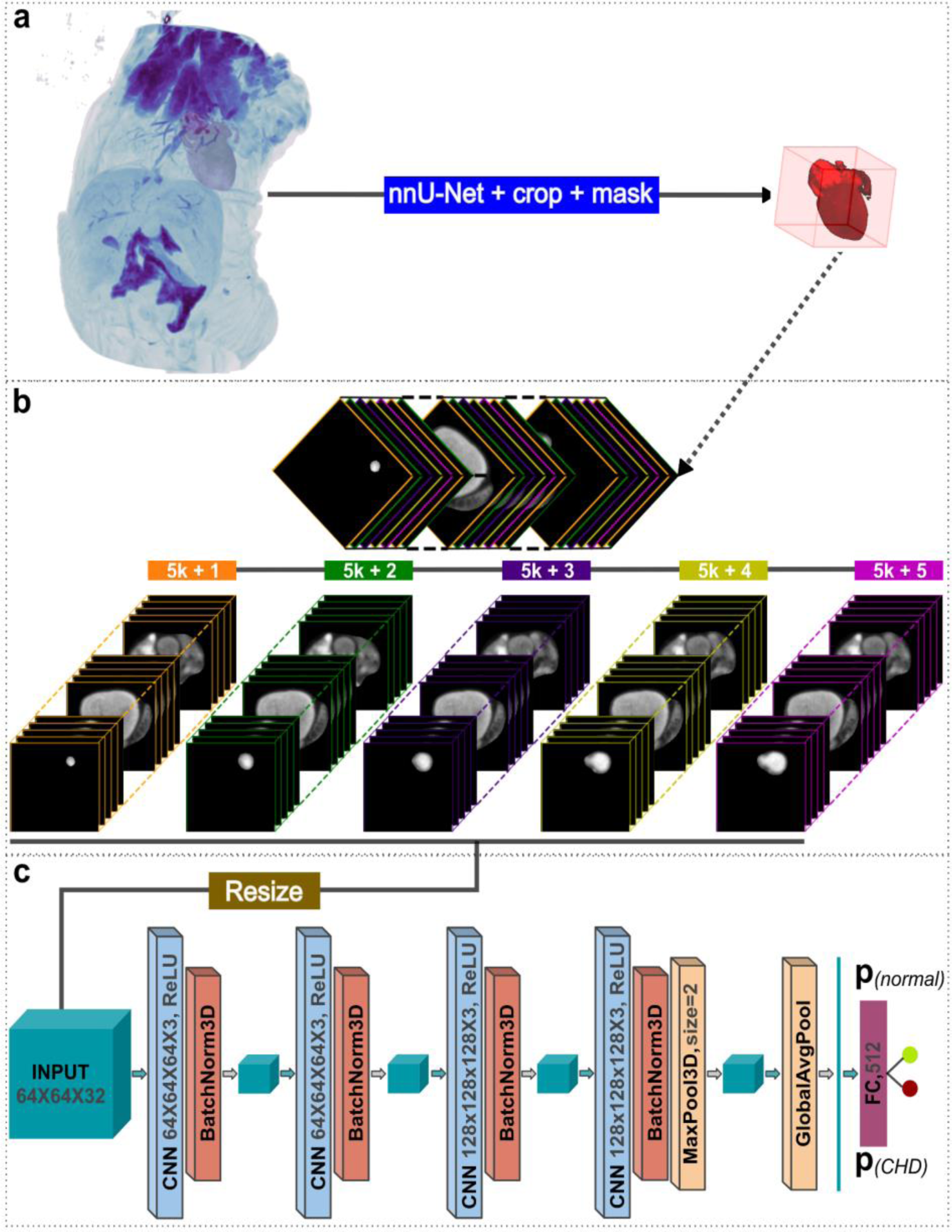
Deep learning-based framework to identify abnormal heart morphology. **a)** A nnU-Net is used to segment hearts from μCT scans. The 3D renderings show the entire μCT scan on the left, and a bounding box around the segmented heart on the right (with 5 voxel padding along the three axes). This bounding box is used to define inputs for the classification module in **c**. **b)** From the stack of slices in this bounding box, we extract five distinct 3D images by sampling every fifth slice, as illustrated (each color corresponds to a distinct 3D images), leading to five times more 3D training images. **c)** The diagnosis module is a 3D CNN that takes 3D images resized to 64 x 64 x 32 voxels as input and comprises four convolutional blocks, a 3D max-pooling layer, a global average pooling layer, a dense (fully connected, FC) layer with 512 neurons and an output layer with two neurons (encoding the predicted probabilities of hearts having CHD or being normal). Each convolutional block comprises one 3D convolutional layer and a 3D batch normalization layer.

Our 3D CNN consisted of four 3D convolutional blocks, one fully connected layer, and an output layer with two neurons (Figure 2c). The first three 3D-CNN blocks were each composed of a 3D-CNN layer followed by a 3D batch normalization layer. The last block consisted of one 3D-CNN layer, a 3D batch normalization layer, and a max pooling layer (pool size = 2). The output of the last convolutional layer underwent a global average pooling, followed by a fully connected layer with 512 neurons, and finally an output layer with two neurons to calculate the probabilities of the heart being normal or having CHD (Figure 2c).

We used the stochastic gradient descent (SGD) optimizer with a learning rate of 0.01 to optimize the model over a maximum of 100 epochs. To address class imbalance, we used a class-weighted cross-entropy loss function *L* = −[*w*_1_ ylog *p* + *w*_0_(1 − *y*) log(1 − *p*)] with weights *w*_0_ = (*n*_0_ + *n*_1_)⁄2*n*_0_, *w*_1_ = (*n*_0_ + *n*_1_)⁄2*n*_1_, where *n*_0_ and *n*_1_ are the numbers of normal and CHD samples, respectively, *y* is the ground truth and *p* the predicted probability. To avoid overfitting, we stopped training when the validation loss stopped improving after 8 epochs

### Prospective and divergent cohorts

To rigorously evaluate the robustness of the classification model, we collected two additional cohorts after training the model on the initial cohort: a ‘prospective’ cohort and a ‘divergent’ cohort. Supplementary Table 1 presents a comprehensive overview of the diverse mouse lines, genotypes, stages and cardioplegia treatments employed in the initial, prospective, and divergent cohorts and Table 2 provides statistics on the two latter cohorts. The prospective cohort consisted of 18 samples (12 CHD and 6 normal) acquired at the E18.5 stage. It comprised a subset of the mouse lines and genotypes of the initial cohort and used the same cardioplegia treatment, but no novel conditions, hence allowing to evaluate the model on new samples.

**Table 2.**
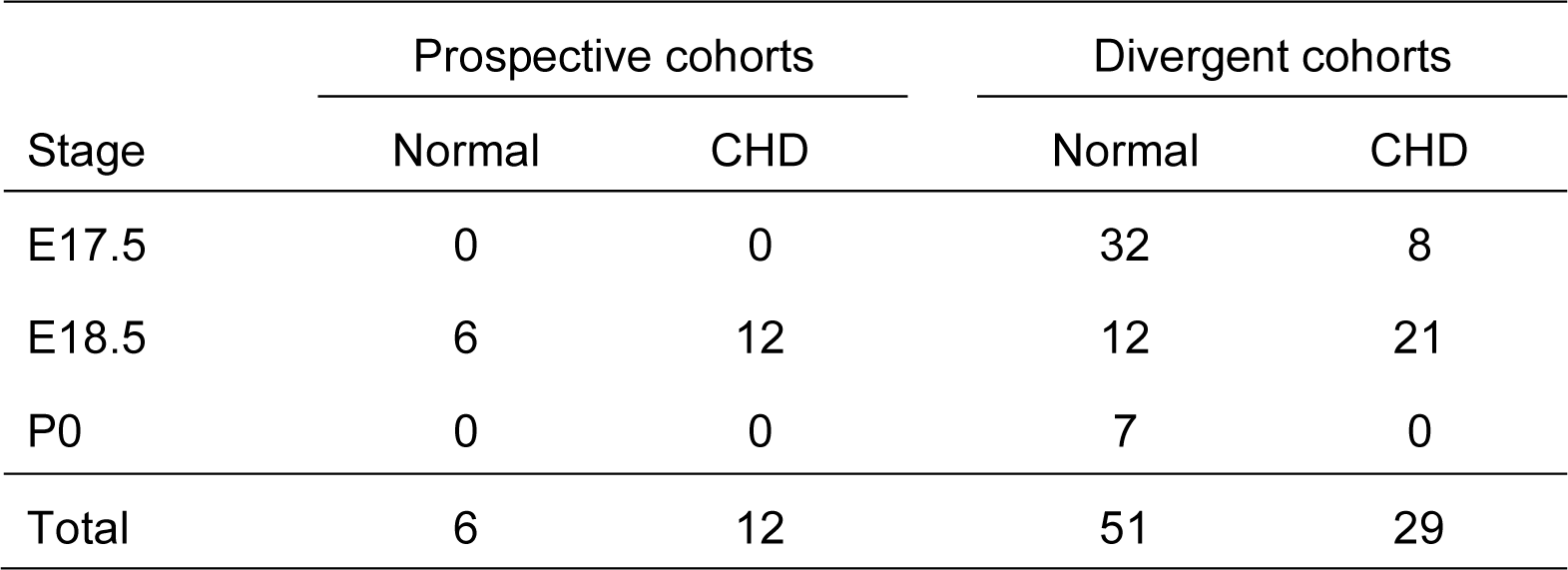
Prospective and divergent cohorts. This table shows the distribution of developmental stages and labels in the prospective and divergent cohorts. CHD: Congenital Heart Defect; E17.5, E18.5, P0: developmental stages at which scans were taken.

The divergent cohort consisted of 80 samples (29 CHD and 51 normal) and introduced entirely novel combinations of mouse lines, genotypes, cardioplegia treatments, and developmental stages, thus offering a stringent test of the model’s robustness. Specifically, we used mouse fetal samples with heterotaxy, including a new line of *Ccdc40^lnks/lnks^* mutants^24^, as well as littermate wild-type controls. *Ccdc40^lnks/lnks^* mutants develop faster, compared to *Nodal* mutants, and were collected the day before birth at E17.5 (equivalent to E18.5 of *Nodal* mutants). *Ccdc40^lnks/lnks^* mutants develop heterotaxy with partial penetrance, as well as situs inversus totalis, while 70% *Ccdc40^lnks/lnks^* mutants have no detectable heart phenotype. Thus, the divergent cohort contains a new phenotype of situs inversus totalis absent from the initial and prospective cohorts. The divergent cohort also contained *Hoxb1^Cre/+^;Nodal^flox/-^* conditional mutants as in the initial cohort but these samples (like all others in the divergent cohort) were treated with a very different cardioplegia method, which is more efficient to relax the cardiac muscle within the endogenous thoracic cavity and thus provides more standardized ventricular wall thickness in controls. Although, the divergent cohort included samples at the E17.5 stage (Figure 5a), which was not represented in the initial and prospective cohorts, E17.5 of the *Ccdc40* mouse line is equivalent to E18.5 of the other lines, because it is one day before birth: indeed we did not detect any significant difference in heart volumes between these two groups (Mann-Whitney U test: n1 = 40, n2 = 107, p = 0.293; Figure 5b).

### Activation map of classification model

In order to highlight features in the μCT images that underlie classification into normal or CHD, we used the Gradient-weighted Class Activation Map (GradCAM) technique^25^. In the resulting activation map, hot colors represent regions that are more important for predicting a specific class (CHD or normal) than regions with cold colors. In our implementation of GradCAM, we first computed feature maps of the last convolutional layer (**A**), then calculated the gradients of the predicted class with respect to these feature maps. Next, these gradients were global-average-pooled to determine the importance weights (*w*). Finally, the GradCAM was computed from the linear combination of feature maps (**A**) and weights (*w*) and application of the ReLu function (i.e. setting all negative values to zero).

### Statistical Analysis

Descriptive statistics of quantitative variables are given as means ± standard deviation (SD) unless otherwise stated. Pixel level segmentation accuracy was quantified using the Dice coefficient, recall, and precision, defined as follows:

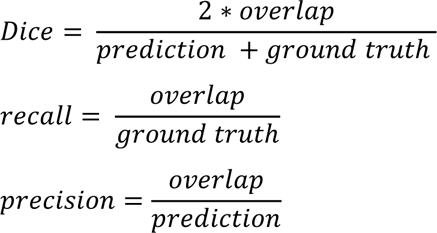

where *prediction* is the number of voxels in the automatically segmented (predicted) 3D masks, *ground truth* is the number of voxels in the manually defined 3D segmentation masks (ground truth), and *overlap* is the number of pixels in the intersections of automated and manual segmentations.

We used the χ2 test to test for dependence between labels (CHD or normal) and stages, and to test for the dependence of segmentation and classification results with labels or stages.

The differences of quantitative variables between two groups were tested by Mann-Whitney U tests. Differences were considered as statistically significant for p-values below 0.05. Statistical analyses were implemented with the *scipy* package (version 1.9.1) within Python 3.9.13.

## Code availability

The MouseCHD Napari plugin was deposited on the Napari Hub at https://www.napari-hub.org/plugins/mousechd-napari and analysis scripts are available at https://github.com/hnguyentt/MouseCHD.

## Results

We implemented and tested a computational pipeline to analyze μCT scans of mouse fetuses, which consists of two main modules: (i) a module for heart segmentation and (ii) a diagnosis module for identification of malformed hearts. The segmentation module extracts heart regions from body μCT scans, while the diagnosis (classification) module predicts the presence or absence of CHD based on segmented hearts. The methods implemented in these modules are described in the Materials and Methods section. We now present an assessment of segmentation and diagnosis results, and of the factors that affect their performance. We first train and test a model on the initial cohort, then evaluate the model’s robustness on the prospective and divergent cohorts, and finally assess the effect of fine-tuning the model on the divergent data. In addition, we present a user-friendly plugin that enables easy application of our segmentation and diagnosis modules, as well as their retraining using other datasets.

### Robust heart segmentation from mouse μCT body scans

The segmentation of the heart serves two purposes: (i) it enables a quantitative analysis of heart volume and shape, and (ii) it facilitates the subsequent identification of malformed hearts by restricting the analyzed images to regions containing the heart.

As described in Materials and Methods, we adopted a deep learning based 3D segmentation framework called nnU-Net^22^, which we trained on 28 manually segmented scans. This framework provides five distinct segmentation models (each corresponding to a distinct data partition) and the final result is obtained by averaging the outputs of the five individual models. Since the heart is a contiguous entity, we further improved the segmentation by keeping only the largest connected component and hence removing fragmented areas.

We evaluated the performance of segmentation in three ways (Figure 3): (i) quantitatively on a subset of the μCT scans that were manually segmented, (ii) semi-quantitatively on the whole dataset, based on a visual evaluation of automated segmentations by an embryologist, and (iii) indirectly, by comparing segmented heart volumes for two different stages.

**Figure 3:**
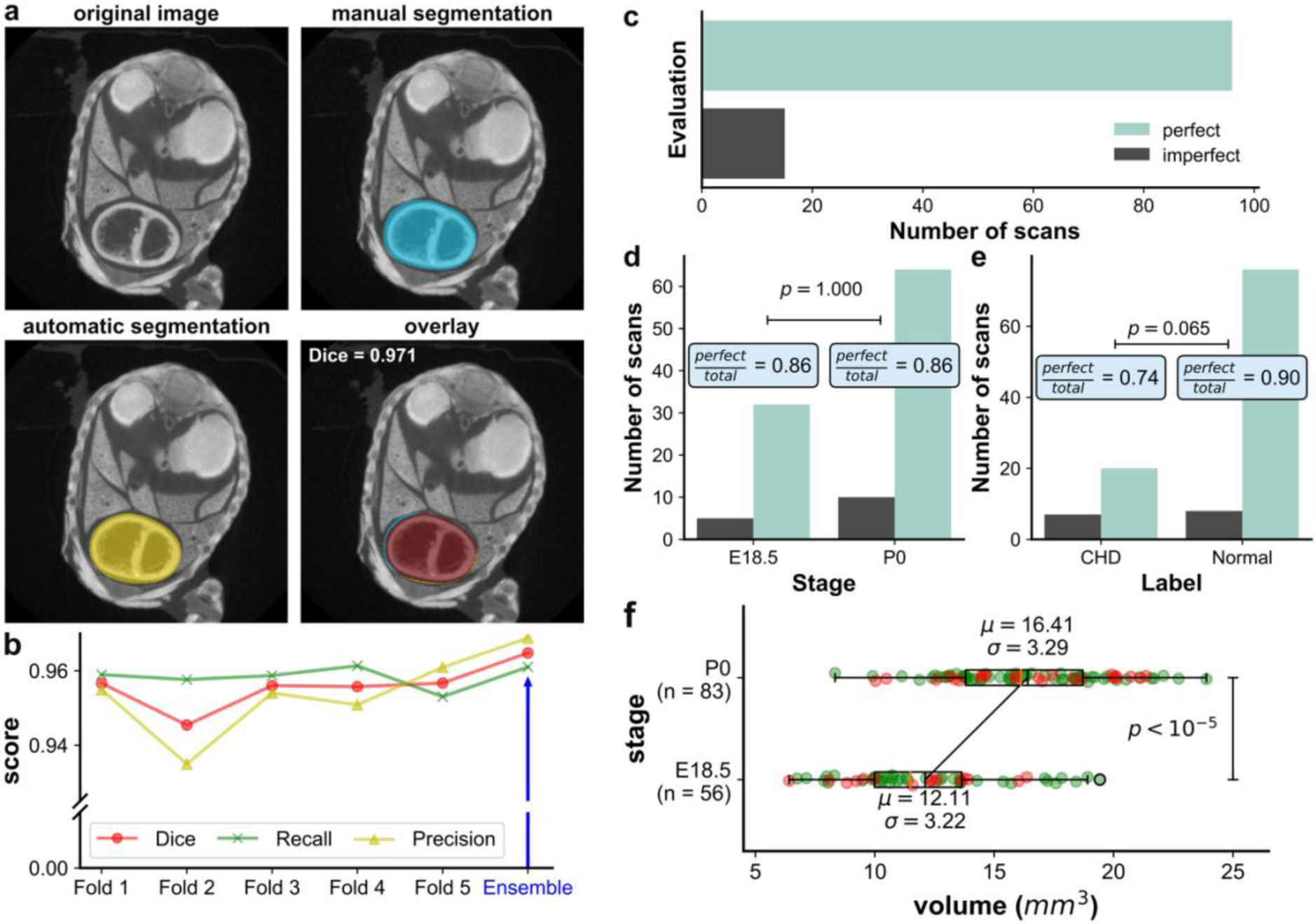
Evaluation of heart segmentation from μCT scans. **a)** Heart segmentation results (evaluated by an expert as perfect), shown on a single slice of a μCT scan: original image (left), manual segmentation (blue), automated segmentation by the nnU-Net (yellow), and overlap between automated and manual segmentations (red). **b)** Segmentation performance metrics for the five models trained by nnU-Net on five data folds. Dice coefficient, recall, and precision are shown. Ensemble designates the pixel-wise average over the five folds. **c)** Horizontal bars show the segmentation performance evaluated by an expert: an embryologist assessed 111 heart segmentations (28 training hearts excluded) visually and classified 96 hearts (86%) as perfectly segmented. **d,e)** Vertical bars show the number of perfectly segmented hearts according to stage (**d**) or label (CHD or normal) (**e**). The proportion of perfectly segmented hearts did not differ significantly according to stage or label (p>0.05 in both cases). **f)** Boxplots compare the heart volumes as measured from 139 automated segmentations for the two developmental stages. Green dots correspond to normal hearts, red dots to hearts with CHD.

First, an embryologist performed manual segmentations of all 2D slices in 12 μCT scans that were not part of the training data, thus providing a ground truth for quantitative assessment of automated segmentations (Figure 3a). We used the Dice coefficient, recall, and precision to measure the agreement between predicted and manual segmentation, each of which would reach 100% for perfect segmentations. The nnU-Net segmentation achieved a Dice coefficient of 96.5%, a recall of 96.1%, and a precision of 95.9%, indicating highly accurate segmentations (Figure 3b).

Second, an embryologist qualitatively assessed the segmentations for the complete dataset of 139 μCT scans and labeled them as either perfectly segmented, over-segmented, or under-segmented (see Materials and Methods). Ignoring the 28 scans used for training, we found that 86% of hearts (n=96) were perfectly segmented, while 14 % (n=15) hearts were under- or over-segmented (Figure 3c). For these 14% of imperfect segmentations, the discrepancies between automated and manual segmentations remained minor (see Figure 3a and Supplementary Figure 1 for examples), and mainly restricted to the great vessels connecting the heart. The segmentation performance, as measured by the percentage of perfectly segmented scans, did not depend on the developmental stage, since it was the same (86%) for both E18.5 and P0 (χ^2^(2, N = 111) = 1.15, p = 1.00) (Figure 3d). Percentages of perfectly segmented scans for normal and CHD hearts were 90% and 74%, respectively, but this difference was not statistically significant (χ^2^(2, N = 111) = 4.72, p = 0.07) (Figure 3e). Thus, the segmentation module exhibits high performance across the entire dataset irrespective of differences in heart morphology.

Third, we used our automatic segmentation to calculate heart volumes in the entire data set. Since the heart grows in size during gestation^26,27^, accurate segmentation should yield an increase in volume between the E18.5 stage and birth (P0). This was indeed the case, since segmented heart volumes at birth (16.41 ± 3.29 mm^3^) were significantly larger than in the E18.5 stage (12.11 ± 3.22 mm^3^) (Mann-Whitney U test: n1 = 56, n2 = 83, p < 10^−5^) (Figure 3f).

These three distinct validations demonstrate high-quality segmentation of mouse hearts from μCT scans regardless of developmental stages and cardiac anomalies.

### Accurate diagnosis of malformed hearts

We next assessed the performance of our 3D-CNN for detecting cardiac malformations (Figure 4). Because of the low number of available μCT scans, especially for hearts with CHD (n=38 out of a total of n=139) we evaluated the model using five-fold cross-validation, whereby the data were split into five stratified folds, allowing to train five models on four folds each, and testing each model on the remaining fold (Table 3). We stratified the data such as to ensure that the ratios of hearts with or without CHD and from the E18.5 or P0 developmental stages were similar between all folds. Visual comparisons of the pixel intensity histograms within and between folds, or in CHD vs normal hearts did not reveal any obvious difference between these groups and comparisons of the images using dimensionality reduction (t-SNE) did not show any correlation that might bias the evaluation (Supplementary Figure 2a-c).

**Figure 4:**
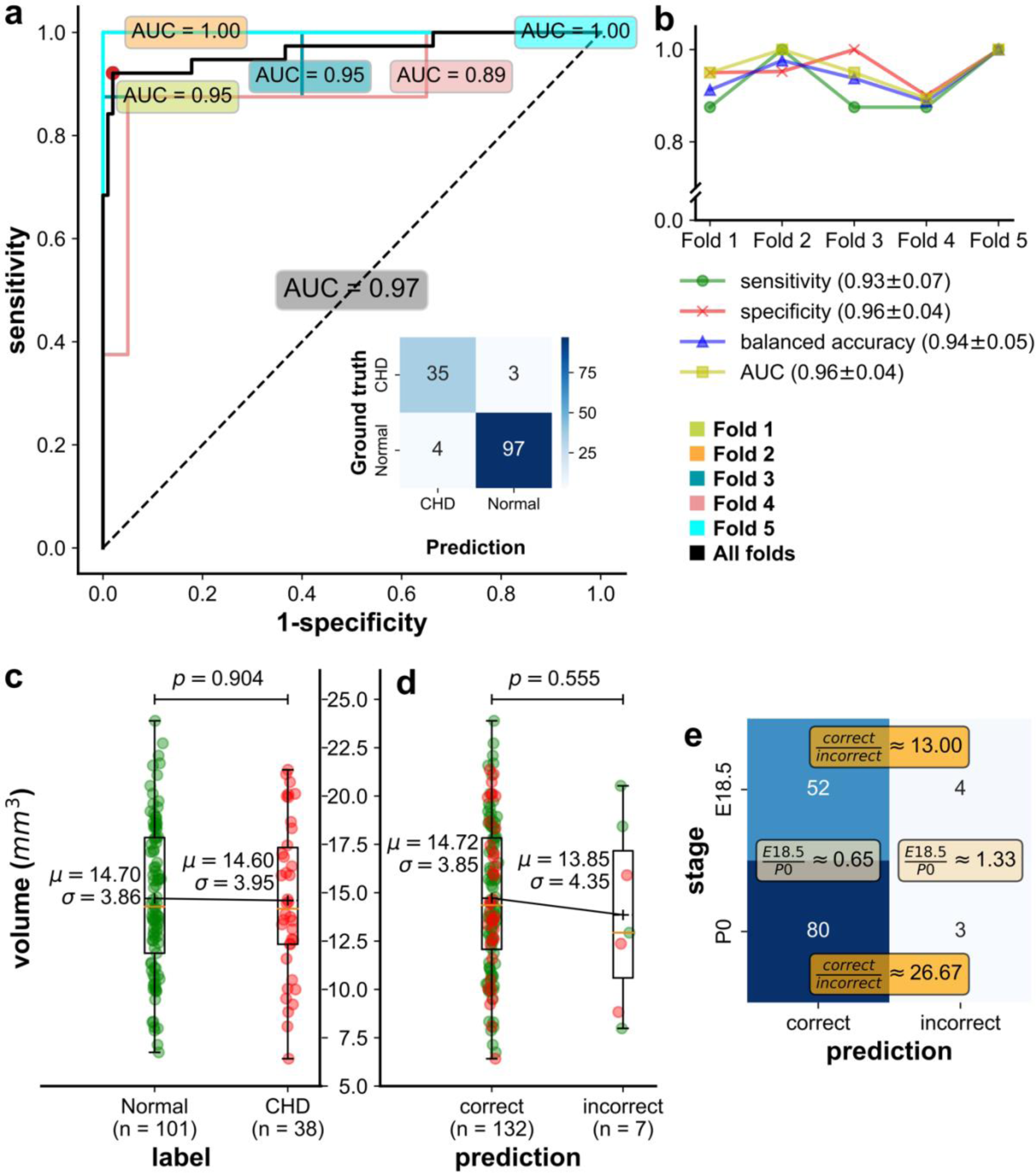
Evaluation of the diagnosis module for mouse fetal hearts. **a)** Diagnostic performance for detection of malformed hearts as measured by the sensitivity vs. specificity trade-off (ROC) curve. The area under the curve (AUC) of five models/folds (each shown by a different color) ranges from 0.89 to 1.00, and the overall AUC (black line) is 0.97. The confusion matrix (counting the number of samples for each predicted class and ground truth class) corresponds to the ensemble of five folds with AUC=0.97. **b)** Different metrics for diagnostic performance in five-fold cross-validation. The detection model achieved an average AUC, sensitivity, specificity, and balanced accuracy of 0.96 ± 0.04, 0.93 ± 0.07, 0.96 ± 0.04, and 0.94 ± 0.05, respectively, as assessed by five-fold cross-validation on 139 hearts. **c, d)** Boxplots comparing the heart volumes for CHD vs normal cases (**c**) and for correct vs. incorrect classifications (**d**). **e)** Contingency table counting the number of samples with correct or incorrect predictions according to the developmental stage. Green dots correspond to normal hearts, red dots to hearts with CHD.

**Table 3.**
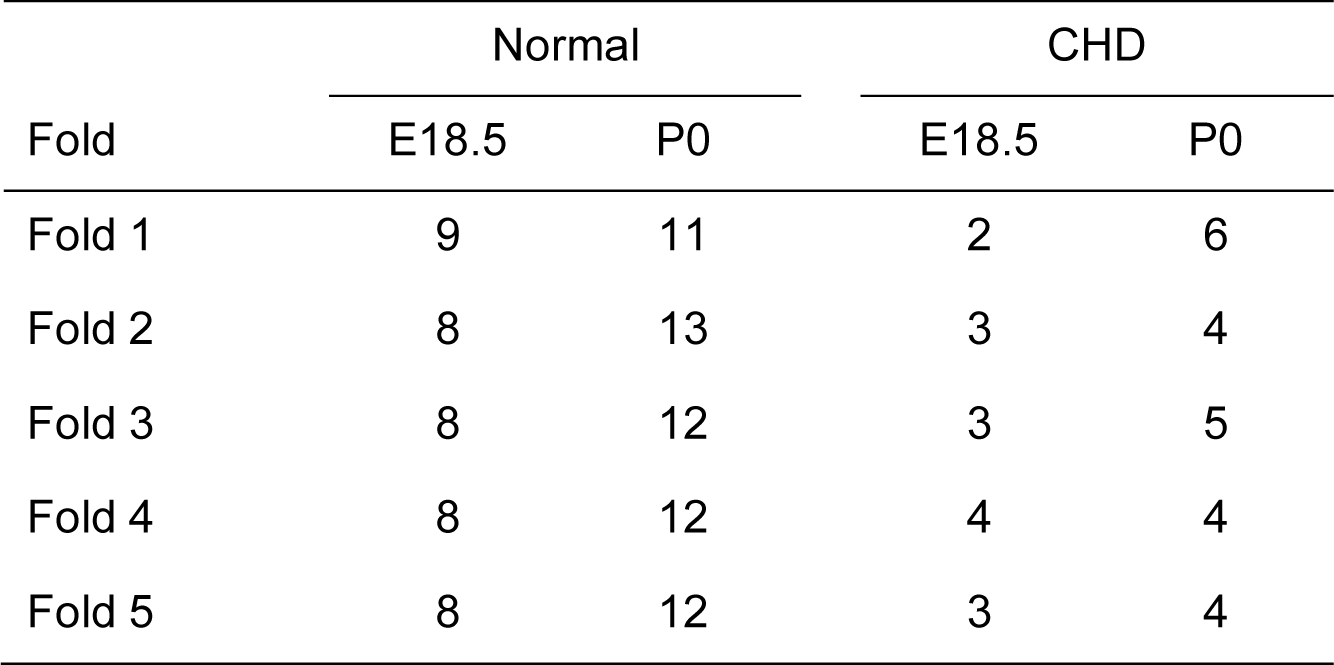
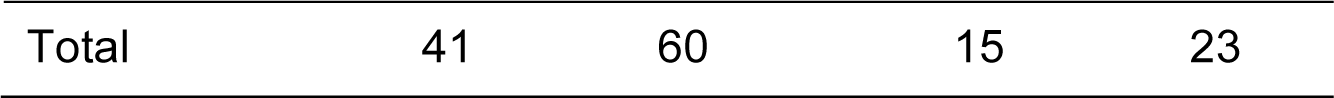
Data split for training and testing the heart anomaly diagnosis module. Table entries indicate the number of scans included in each of the five folds for cross-validation. All samples are from the initial cohort. CHD: Congenital Heart Defect; E18.5, P0: developmental stages at which scans were taken.

Detection implies a tradeoff between sensitivity and specificity, both of which depend on the threshold chosen for binary classification (CHD vs normal). Figure 4a shows the sensitivity vs. specificity for varying thresholds (receiver operating characteristic, or ROC, curve) for each of the five models/folds (colored curves) as well as the average ROC curve (black). Area under the curve (AUC), a measure of classification accuracy that equals 1.0 for perfect diagnosis, ranged from 0.89 to 1.00 for the five models, with an average of 0.96. At the chosen threshold (p=0.5), the average sensitivity was 93%, the average specificity was 96%, and the class-balanced accuracy was 94% (Figure 4b), indicating good classification performance.

For comparison, we also trained and tested two other diagnosis models on the entire μCT scans without using the heart segmentation (Supplementary Figure 3). The first model was trained using the same 3D-CNN architecture as above, but on the body scans rather than segmented heart regions. This model only achieved a sensitivity of 63%, a specificity of 90%, a balanced accuracy of 77%, and an AUC of 0.77 (Supplementary Figure 3a). The second model was trained with attention-based deep multiple instance learning (AttMIL) ^28^ on heart slices, using an architecture shown in Supplementary Figure 3c. This AttMIL model achieved a sensitivity of 78%, a specificity of 95%, a balanced accuracy of 86.5%, and an AUC of 0.92 (Supplementary Figure 3a). Thus in both cases, classification performance was lower than with segmentation, likely because the model learned confounding factors located outside the heart rather than features intrinsic to the hearts, which make up only ∼10% in volume of the whole μCT scans (Supplementary Figure 3b,d). This shows the importance of locating the heart and the benefit of performing heart segmentation for the diagnosis pipeline.

Among potential confounders of disease diagnosis are heart volumes and developmental stage. However, heart volumes did not differ significantly between scans with and without CHD (Figure 4c; Mann-Whitney U-test p=0.90) or between correctly and incorrectly classified scans (Figure 4d; Mann-Whitney U-test p=0.56). Likewise, the proportion of correct classifications did not significantly differ between E18.5 and P0 hearts (χ2(1, N = 139) = 0.29, p = 0.59) (Figure 4e). Thus, our model appears to have learned to identify heart malformations irrespective of heart size or developmental stage.

In the following, based on the results of our cross-validation analysis, we consider a single ‘final’ model trained on the entire data set of N = 139 samples from the initial cohort.

### Robust diagnosis on prospective and divergent cohorts

To assess our model’s performance in real world settings, we next tested it on samples obtained exclusively after model training (Figure 5). We started with the prospective cohort, which as mentioned above includes a subset of the conditions contained in the initial cohort (see Supplementary Table 1). The distribution of cardiac phenotypes in this data set did not significantly differ from the initial cohort used for model training (Supplementary Figure 4a,b). Evaluation of our model on this data set again showed excellent performance, with a balanced accuracy of 96%, a sensitivity of 92%, and perfect specificity and AUC of 100% (Figure 5c,d). These findings validate the model’s adaptability to novel data in absence of changes in mouse lines, developmental stages, genotypes or cardioplegia treatment.

**Figure 5:**
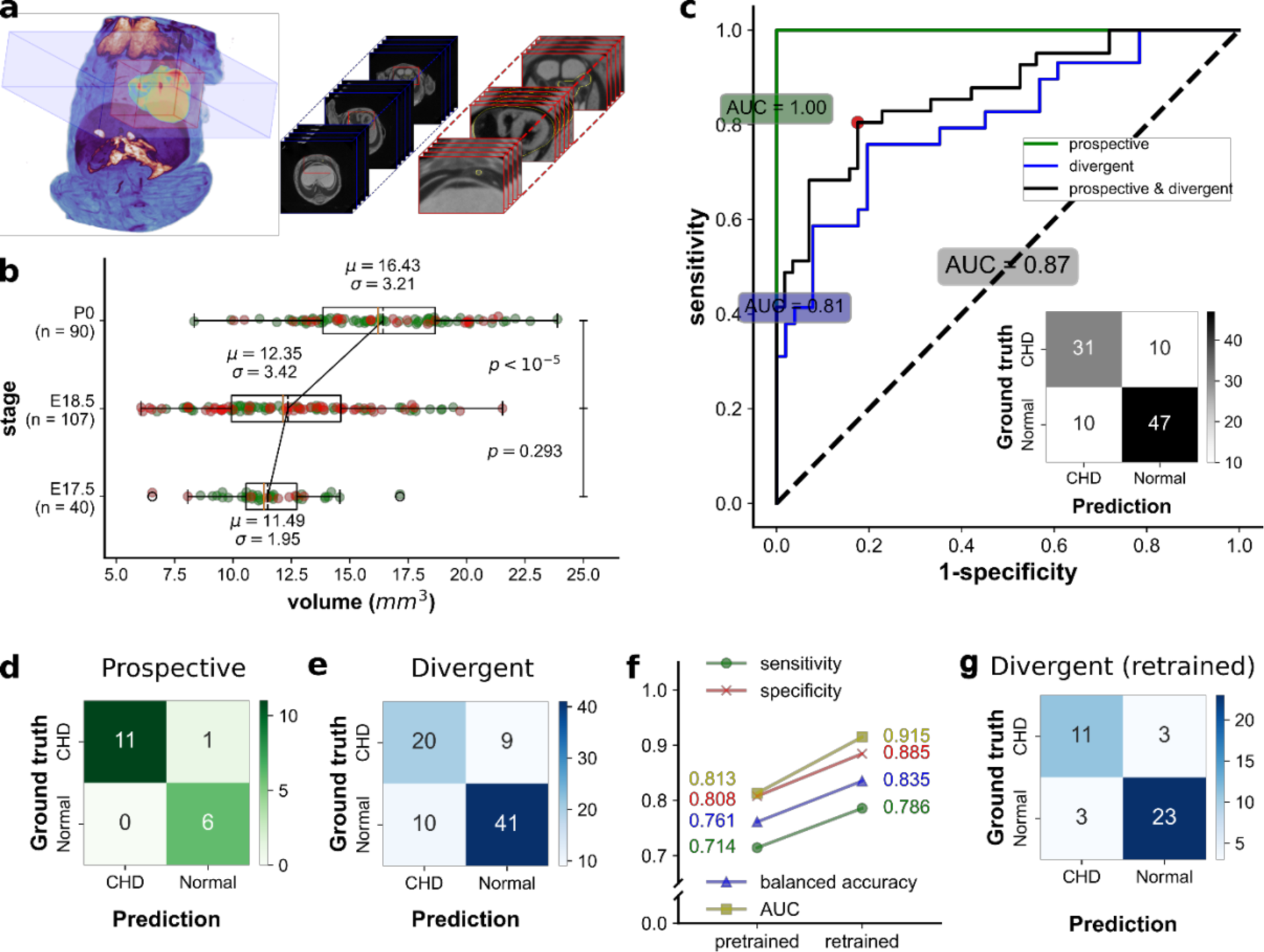
Evaluation of the diagnosis module on prospective and divergent cohorts. **a)** Body μCT scan of a mouse fetus at 17.5 days of development (E17.5) from the *Ccdc40* line, shown as 3D rendering (left) and stacks of 2D slices (right). Blue bounding box shows the set of 2D thoracic slices containing the heart. The red bounding box is framed around the heart. The corresponding 2D image stacks are shown with matching color frames. **b)** Boxplots compare the heart volumes at different stages (E17.5, E18.5, and P0). Heart volumes differ significantly between stages P0 and E18.5 (p < 10^-5^), but not between E17.5 (*Ccdc40* line) and E18.5 (all other lines) (p = 0.293), because they both correspond to one day before birth. **c)** Diagnostic performance of the model trained on the initial cohort for detection of malformed hearts, assessed by the sensitivity vs. specificity trade-off (ROC) curve. The area under the curve (AUC) for prospective data (green) and divergent data (blue) is 1.0 and 0.81, respectively, and the overall AUC (prospective and divergent data, black line) is 0.87. The confusion matrix shows the counts of samples for each predicted and ground truth class, for the combined prospective and divergent data. **d, e)** Confusion matrices of the model trained on the initial cohort using test data from the prospective cohort (d) and the divergent cohort (e). **f)** Performance of the model on 40 divergent test data before (left) and after (right) finetuning on distinct data from the divergent cohort. Sensitivity increases from 0.714 to 0.786, specificity from 0.808 to 0.885, balanced accuracy from 0.761 to 0.835, and AUC from 0.813 to 0.915. **g)** Confusion matrix of the model finetuned on the prospective cohort and a subset of the divergent cohort, and tesed on the remaining subset of the divergent cohort.

For a more stringent test, we next turned to the divergent cohort, which unlike the prospective cohort contains many differences relative to the training data. These include the presence of different mouse lines, and application of a very different cardioplegia treatment (Supplementary Table 1). Importantly, the distribution of phenotypes in this data set was significantly different (χ2 test: p=0.03), with the inclusion of novel phenotypes -in particular situs inversus totalis-that were absent from the initial cohort and hence unseen by the model (Supplementary Figure 4a,b). Despite all these differences with the training data, our model achieved relatively good performance (balanced accuracy: 75%, sensitivity: 69%, specificity: 80%, AUC: 81%) (Figure 5c,e). This suggests that the model has captured image features that robustly distinguish normal from abnormal hearts.

### Finetuning improves performance on the divergent cohort

While these results demonstrate robustness of our model, the lower performance on the divergent cohort compared to the initial and the prospective cohorts calls for an improved model. Because of the many differences between the divergent and the initial cohort mentioned above, improved performance on the divergent cohort can be expected by finetuning the model to account for these differences. In order to avoid any bias in the testing data, we divided the divergent cohort evenly into two subsets, ensuring similar distributions of genotypes, mouse lines, cardioplegia treatment, phenotypes, and predictions from the pretrained model (see Supplementary Table 2 for details). We then finetuned our model (pretrained on the initial cohort) on one subset of the divergent cohort, combined with the entirety of the prospective cohort, and tested it on the remaining divergent cohort subset.

Finetuning led to improved performance on all metrics: sensitivity increased from 71% to 79%, specificity from 81% to 88%, and the AUC jumped from 81% to 91% (Figure 5f,g). These results demonstrate the versatility of our tool, as it can be easily retrained to bolster diagnostic performance for diverse data sets.

### Easy-to-use and retrainable Napari plugin to visualize and analyse heart defects

In order to empower embryologists and others to reutilize and improve our pipeline without requiring coding skills, we built them into a user-friendly plugin – MouseCHD-for Napari, a widely used and open source visualization and image analysis platform^19^ (Figure 6). MouseCHD is available on both the Napari plugin repository and GitHub, along with sample data (https://github.com/hnguyentt/mousechd-napari). After importing body μCT scans with a NifTI^29^, DICOM^30^ or Nrrd^31^ format, users can choose between three different tasks: (i) segmentation of the heart, (ii) diagnosis of malformation, and (iii) retraining of the diagnosis model (Figure 6a,b). The output of the segmentation module is a 3D mask of the heart (Figure 6c), while the output of the diagnosis module is a predicted probability for the heart to be normal or to have CHD. Thanks to Napari’s built-in features, users can easily visualize images in 2D or 3D and interact with them, e.g. by rotating or adjusting contrast and transparency (Figure 6d). This facilitates a thorough inspection of heart morphology and segmentation results. (Note that the heart segmentation module requires at least one graphics processing unit (GPU). For users who do not possess local GPUs, we provide instructions on how to run the plugin using remote high-performance computing clusters or cloud services. Detailed guidelines are available at: https://github.com/hnguyentt/mousechd-napari/blob/master/docs/server_setup.md)

**Figure 6:**
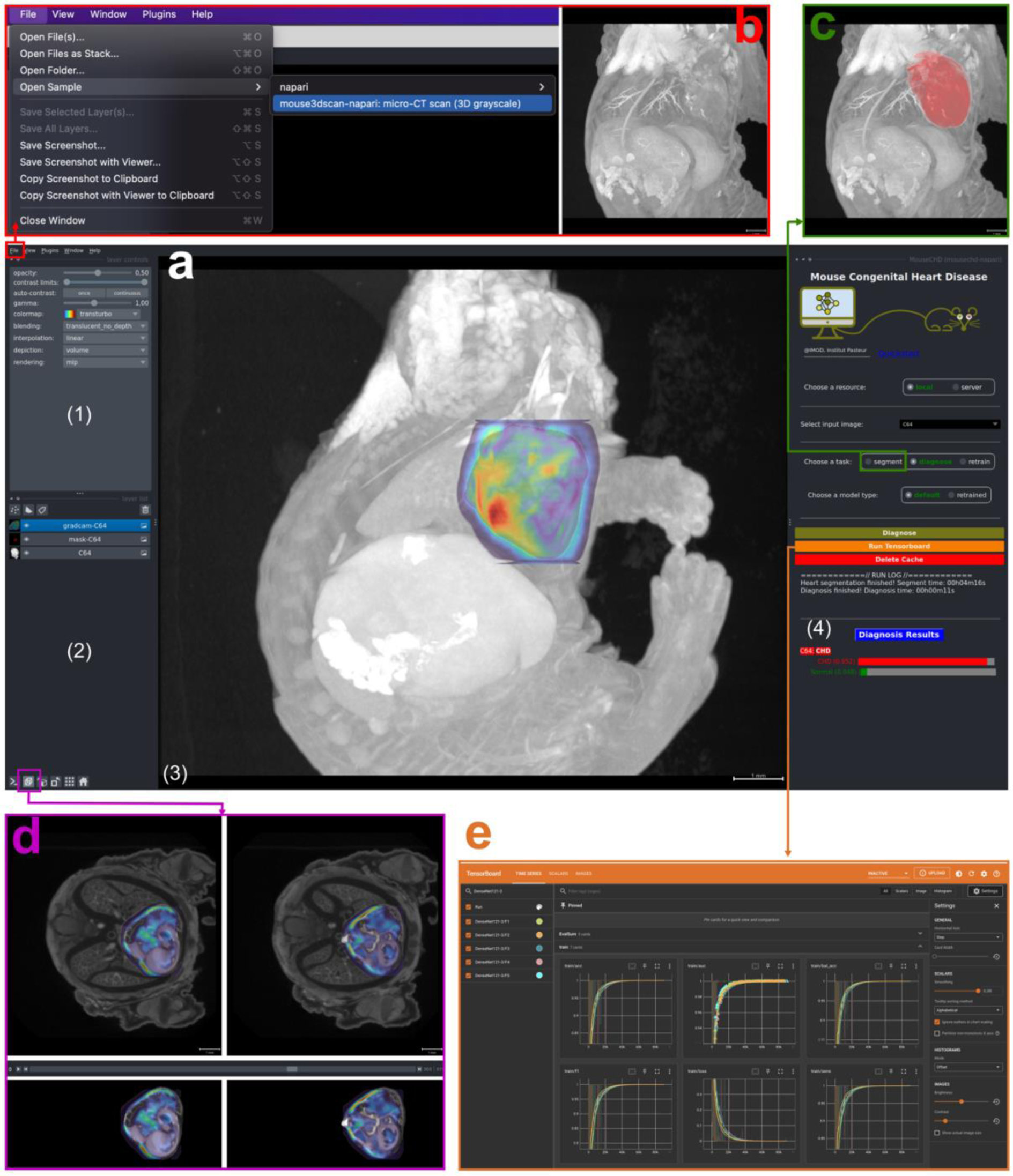
Napari plugin for heart segmentation and diagnosis of heart malformation. **a)** Plugin screenshot. The Napari graphical user interface is divided in four panels: (1) layer control to adjust visualization parameters such as opacity, contrast, and color maps, for each layer; (2) list of all visualization layers, which can be individually displayed or hidden using the eye icon; (3) canvas for image visualization, (4) the “MouseCHD” plugin panel lets users choose between three tasks: heart segmentation, CHD diagnosis, or re-training of the diagnostic model. The diagnosis result is shown on the right as two probabilities for CHD (red) and normal (green). The canvas (3) displays the input image, the heart segmentation, and the GradCAM activation map, as three distinct layers in 3D. Users can rotate the image in 3D. **b)** Sample image, which can be loaded by choosing “File” → “Open Sample” → “mouse3dscan-napari: micro-CT scan (3D grayscale)”. **c)** Heart segmentation result displayed in 3D in the canvas (3) when the user chooses the segmentation task. **d**) 2D views: users can click on the viewer button (cube icon in the magenta box) and use the slider to select 2D image slices. **e)** Re-training: the user provides the data folder containing CHD and normal images organized into two sub-folders: “CHD” and “Normal”. The plugin automatically preprocesses the data, segments hearts and re-trains the diagnosis (classification) model. The plugin allows users to monitor the training process through dynamic visualization of key performance metrics – such as accuracy, recall and precision – plotted against training epochs using TensorBoard. Users can launch TensorBoard by clicking the “Run TensorBoard” button and following the provided link.

In addition, we implemented a visualization technique called gradient-weighted class activation mapping (GradCAM) ^25^), that highlights the image regions that most strongly contribute to the classification output (Figure 6a,d). Preliminary inspection indicates that for true positive CHD detections, GradCAM highlights regions between ventricles and atria in 30 out of 35 true positive cases (86%) (Supplementary Figure 5). Since atrioventricular septal defects are a common feature of all CHD cases in our initial data set (Figure 1b), the model seems to correctly identify these defects for CHD prediction. In addition, GradCAM visualizations can also guide further improvements to the classification model itself. For example, when we applied the AttMIL model to the heart slices, GradCAM tended to highlight regions inside the mouse lungs, instead of the heart (Supplementary Figure 3d). This is relevant to our dataset of samples, because the heterotaxy syndrome has associated lung defects (isomerism) with full penetrance in *Nodal* mutants^5,20^. Yet, lung defects cannot be taken as predictive of cardiac defects. Such observations led us to introduce the heart segmentation module, which proved essential to achieve satisfactory classification performance based on cardiac features only (see above).

Most importantly, while MouseCHD provides direct access to our best-performing diagnosis model described above, it also enables users to retrain the model on their own data, potentially using categories of CHD not present in our dataset (Figure 6e). This retraining capability comes with tools to monitor training progress and will make it easy for embryologists to generate tailored diagnosis models that are adapted to -and likely outperform the model provided by us-on their own specific data.

Thus, MouseCHD offers researchers and clinicians in cardiology a number of ready-to-use analysis and visualization tools for studying heart morphology and anomalies in mouse fetuses.

## Discussion

We presented a fully automated pipeline to diagnose heart malformations in mouse fetuses using 3D µCT-scans. Our pipeline includes deep learning-based segmentation of the heart and a 3D-CNN model to detect heart structural anomalies. We provide the pipeline as a user-friendly tool within Napari that will enable researchers to quickly and easily analyze 3D images and identify cardiac defects.

The automated heart segmentation achieved a Dice score of 96% when compared with manual segmentations by a human expert. While implemented to facilitate subsequent diagnosis, this segmentation tool is time-saving and already enables visual insights into heart morphology and quantitative measurements such as the total heart volume.

Despite having been trained on a fairly limited amount of µCT scans (∼80 normal and ∼30 CHD cases), our diagnostic module detected CHD with a sensitivity, specificity, and AUC of 92%, 96%, and 97% respectively in the initial cohort. The model performed even better (sensitivity: 92%, specificity: 100%, AUC: 100%) on a small prospective cohort acquired later and containing samples corresponding to a subset of the conditions in initial cohort. The balanced accuracy (95% and 96% for the initial and prospective cohorts) was on par with that reported in related studies on human imaging data of the heart, despite the much larger size of human organs and the use of very different imaging techniques^32,33^. When tested on a divergent cohort containing differences in genotypes, phenotypes and cardioplegia treatment, our model still performed well, and much better than chance (sensitivity: 71%, specificity: 81%, AUC: 81%), indicating a reasonable degree of robustness of the learned features to these differences. Importantly, finetuning the model on a subset of the divergent cohort in a transfer learning approach^39^ led to markedly improved performance (sensitivity: 79%, specificity: 88%, AUC: 91%), underscoring our model’s adaptability to handling new data distributions and its utility for practical applications. Finally, our method includes an implementation of Grad-CAM, a tool to visually highlight the image regions that underly diagnostic of CHD. Although more work is needed to validate this tool’s ability to directly highlight cardiac malformations, it may serve as a basis for visual explanations of CHD diagnosis by deep learning in the future.

In line with the growing importance of deep learning in medical image analysis and diagnosis of human cardiac conditions^34^, including CHD^35^, our approach uses CNNs that automatically learn complex and highly predictive features. This approach stands in contrast with previous studies that use handcrafted features^12,14,15^, for automated phenotyping of mouse fetuses from µCT data. The striking performance of deep learning-based 2D image classification relies strongly on the availability of large annotated image data bases, such as ImageNet^36^, which includes more than 14 million 2D images scraped from the Internet. By contrast, data bases of 3D medical images, such as The Cancer Imaging Archive (TCIA^37^), currently only contain thousands of images. As a consequence, there is still a dearth of easily accessible pretrained 3D classification models for medical imaging data in general and for cardiology in particular. Our open-access, user-friendly and -crucially-retrainable diagnosis plugin provides biologists with a flexible tool to automate a personalised classification of their own cohorts. This could stimulate the growth of annotated 3D-cardiac image data bases and thereby allow to train models with further improved diagnostic accuracy.

A limitation of the present study is its reliance on a single mouse disease (heterotaxy), developmental window and imaging technique. While our analysis of a divergent cohort indicates strong transferability of the learned features it also showed reduced performance in absence of retraining. Therefore, an obvious follow-up is to increase the diversity of the training data, for example using external cohorts such as IMPC^38^. A model pretrained on a larger diverse data set (e.g. IMPC) can then still benefit from fine-tuning on local data (such as the µCT-scans used in this study) to further improve diagnostic accuracy. Our Napari plugin will facilitate such retraining. Beyond the supervised learning approaches with 3D-CNNs implemented here, follow-up work may also benefit from recent advances such as vision transformers or self-supervised methods that use pre-text tasks to extract relevant features from related unannotated data such as publicly available 3D µCT-scan images from unrelated cohorts^40^. Additionally, more elaborate pre-processing may improve the performance and robustness of the model as suggested by previous studies that emphasized the importance of spatial registration^12^ and image normalization to address varying voxel intensities across different µCT imaging settings^14^. Finally, it will be important to extend this study towards distinguishing between different types of cardiac defects. The methods and open software tools described here are expected to stimulate such developments. Even without awaiting such improvements, we believe that our method’s ability to automate the analysis of cardiac morphology in mice will accelerate the identification of mechanisms of congenital heart defects.

## Acknowledgements

We thank Arthur Herbout for his work in a related effort preceding this study and Cédric Thépenier for insightful comments on the manuscript. This work was funded by the Inception project through Investissement d’Avenir grant ANR-16-CONV-0005 and in part by Institut National du Cancer (Programme de recherche translationnelle en cancérologie 2020/ PRT-K 2020). The Meilhac laboratory is supported by core funding from the Institut Pasteur and state funding from the Agence Nationale de la Recherche under the “Investissements d’avenir” program (ANR-10-IAHU-01). AOS is supported by the Pasteur - Paris University (PPU) International PhD Program, SMM and AD are INSERM research scientists.

## References

1. Hoyert, D. & Gregory C.W., E. Cause-of-Death Data From the Fetal Death File, 2018–2020. https://stacks.cdc.gov/view/cdc/120533 (2022) doi:10.15620/cdc:120533.

2. Ely, D. & Driscoll, A. Infant mortality in the United States, 2020: Data from the period linked Birth/infant death file. Natl. Vital Stat. Rep. 71, 18 (2022).

3. Houyel, L. & Meilhac, S. M. Heart Development and Congenital Structural Heart Defects. Annu. Rev. Genomics Hum. Genet. 22, 257–284 (2021).

4. Diab, N. S. et al. Molecular Genetics and Complex Inheritance of Congenital Heart Disease. Genes 12, 1020 (2021).

5. Desgrange, A., Lokmer, J., Marchiol, C., Houyel, L. & Meilhac, S. M. Standardised imaging pipeline for phenotyping mouse laterality defects and associated heart malformations, at multiple scales and multiple stages. Dis. Model. Mech. 12, dmm038356 (2019).

6. MGI-Mouse Phenotypes, Alleles & Disease Models. http://www.informatics.jax.org/phenotypes.shtml.

7. Home. IMPC | International Mouse Phenotyping Consortium https://www.mousephenotype.org/.

8. Hsu, C.-W. et al. Three-dimensional microCT imaging of mouse development from early post-implantation to early postnatal stages. Dev. Biol. 419, 229–236 (2016).

9. Wasserthal, J., et al. TotalSegmentator: robust segmentation of 104 anatomical structures in CT images. Preprint at 10.48550/arXiv.2208.05868 (2022).

10. McGrath, H. et al. Manual segmentation versus semi-automated segmentation for quantifying vestibular schwannoma volume on MRI. Int. J. Comput. Assist. Radiol. Surg. 15, 1445–1455 (2020).

11. Men, K. et al. Fully automatic and robust segmentation of the clinical target volume for radiotherapy of breast cancer using big data and deep learning. Phys. Med. 50, 13–19 (2018).

12. Wong, M. D., Maezawa, Y., Lerch, J. P. & Henkelman, R. M. Automated pipeline for anatomical phenotyping of mouse embryos using micro-CT. Development 141, 2533– 2541 (2014).

13. Chu, Q. et al. CACCT: An Automated Tool of Detecting Complicated Cardiac Malformations in Mouse Models. Adv. Sci. 7, 1903592 (2020).

14. Horner, N. R., et al. LAMA: automated image analysis for the developmental phenotyping of mouse embryos. Development 148, dev192955 (2021).

15. Dickinson, M. E. et al. High-throughput discovery of novel developmental phenotypes. Nature 537, 508–514 (2016).

16. Zouagui, T., Chereul, E., Janier, M. & Odet, C. 3D MRI heart segmentation of mouse embryos. Comput. Biol. Med. 40, 64–74 (2010).

17. LeCun, Y., Bengio, Y. & Laboratories, T. B. Convolutional Networks for Images, Speech, and Time-Series.

18. Li, Z., Liu, F., Yang, W., Peng, S. & Zhou, J. A Survey of Convolutional Neural Networks: Analysis, Applications, and Prospects. IEEE Trans. Neural Netw. Learn. Syst. 33, 6999– 7019 (2022).

19. Sofroniew, Nicholas et al. napari: a multi-dimensional image viewer for Python. Zenodo 10.5281/ZENODO.3555620 (2022).

20. Desgrange, A., Le Garrec, J.-F., Bernheim, S., Bønnelykke, T. H. & Meilhac, S. M. Transient Nodal Signaling in Left Precursors Coordinates Opposed Asymmetries Shaping the Heart Loop. Dev. Cell 55, 413–431.e6 (2020).

21. Vierkotten, J., Dildrop, R., Peters, T., Wang, B. & Rüther, U. Ftm is a novel basal body protein of cilia involved in Shh signalling. Development 134, 2569–2577 (2007).

22. Isensee, F., Jaeger, P. F., Kohl, S. A. A., Petersen, J. & Maier-Hein, K. H. nnU-Net: a self-configuring method for deep learning-based biomedical image segmentation. Nat. Methods 18, 203–211 (2021).

23. Ronneberger, O., Fischer, P. & Brox, T. U-Net: Convolutional Networks for Biomedical Image Segmentation. Preprint at http://arxiv.org/abs/1505.04597 (2015).

24. Becker-Heck, A. et al. The coiled-coil domain containing protein CCDC40 is essential for motile cilia function and left-right axis formation. Nat. Genet. 43, 79–84 (2011).

25. Selvaraju, R. R. et al. Grad-CAM: Visual Explanations from Deep Networks via Gradient-Based Localization. Int. J. Comput. Vis. 128, 336–359 (2020).

26. de Boer, B. A., van den Berg, G., de Boer, P. A. J., Moorman, A. F. M. & Ruijter, J. M. Growth of the developing mouse heart: An interactive qualitative and quantitative 3D atlas. Dev. Biol. 368, 203–213 (2012).

27. Leu, M., Ehler, E. & Perriard, J.-C. Characterisation of postnatal growth of the murine heart. Anat. Embryol. (Berl.) 204, 217–224 (2001).

28. Ilse, M., Tomczak, J. M. & Welling, M. Attention-based Deep Multiple Instance Learning. ArXiv180204712 Cs Stat (2018).

29. mjenkinson. NIfTI-1 Data Format — Neuroimaging Informatics Technology Initiative. https://nifti.nimh.nih.gov/nifti-1/.

30. Digital Imaging and Communications in Medicine (DICOM). (Springer Berlin Heidelberg, Berlin, Heidelberg, 2008). doi:10.1007/978-3-540-74571-6.

31. Teem: nrrd: Definition of NRRD File Format. https://teem.sourceforge.net/nrrd/format.html.

32. Lo, J., Lim, A., Wagner, M. W., Ertl-Wagner, B. & Sussman, D. Fetal Organ Anomaly Classification Network for Identifying Organ Anomalies in Fetal MRI. Front. Artif. Intell. 5, 832485 (2022).

33. Qiao, S. et al. RLDS: An explainable residual learning diagnosis system for fetal congenital heart disease. Future Gener. Comput. Syst. 128, 205–218 (2022).

34. Ahsan, M. M. & Siddique, Z. Machine learning-based heart disease diagnosis: A systematic literature review. Artif. Intell. Med. 128, 102289 (2022).

35. Hoodbhoy, Z. et al. Diagnostic Accuracy of Machine Learning Models to Identify Congenital Heart Disease: A Meta-Analysis. Front. Artif. Intell. 4, (2021).

36. Deng, J. et al. ImageNet: A large-scale hierarchical image database. in 2009 IEEE Conference on Computer Vision and Pattern Recognition 248–255 (2009). doi:10.1109/CVPR.2009.5206848.

37. Clark, K. et al. The Cancer Imaging Archive (TCIA): Maintaining and Operating a Public Information Repository. J. Digit. Imaging 26, 1045–1057 (2013).

38. Muñoz-Fuentes, V. et al. The International Mouse Phenotyping Consortium (IMPC): a functional catalogue of the mammalian genome that informs conservation. Conserv. Genet. 19, 995–1005 (2018).

39. Cheplygina, V., de Bruijne, M. & Pluim, J. P. W. Not-so-supervised: A survey of semi-supervised, multi-instance, and transfer learning in medical image analysis. Med. Image Anal. 54, 280–296 (2019).

40. Tang, Y., et al. Self-Supervised Pre-Training of Swin Transformers for 3D Medical Image Analysis. Preprint at 10.48550/arXiv.2111.14791 (2022).

