## Supplement for "Deep learning-based detection of murine congenital heart defects from µCT scans"

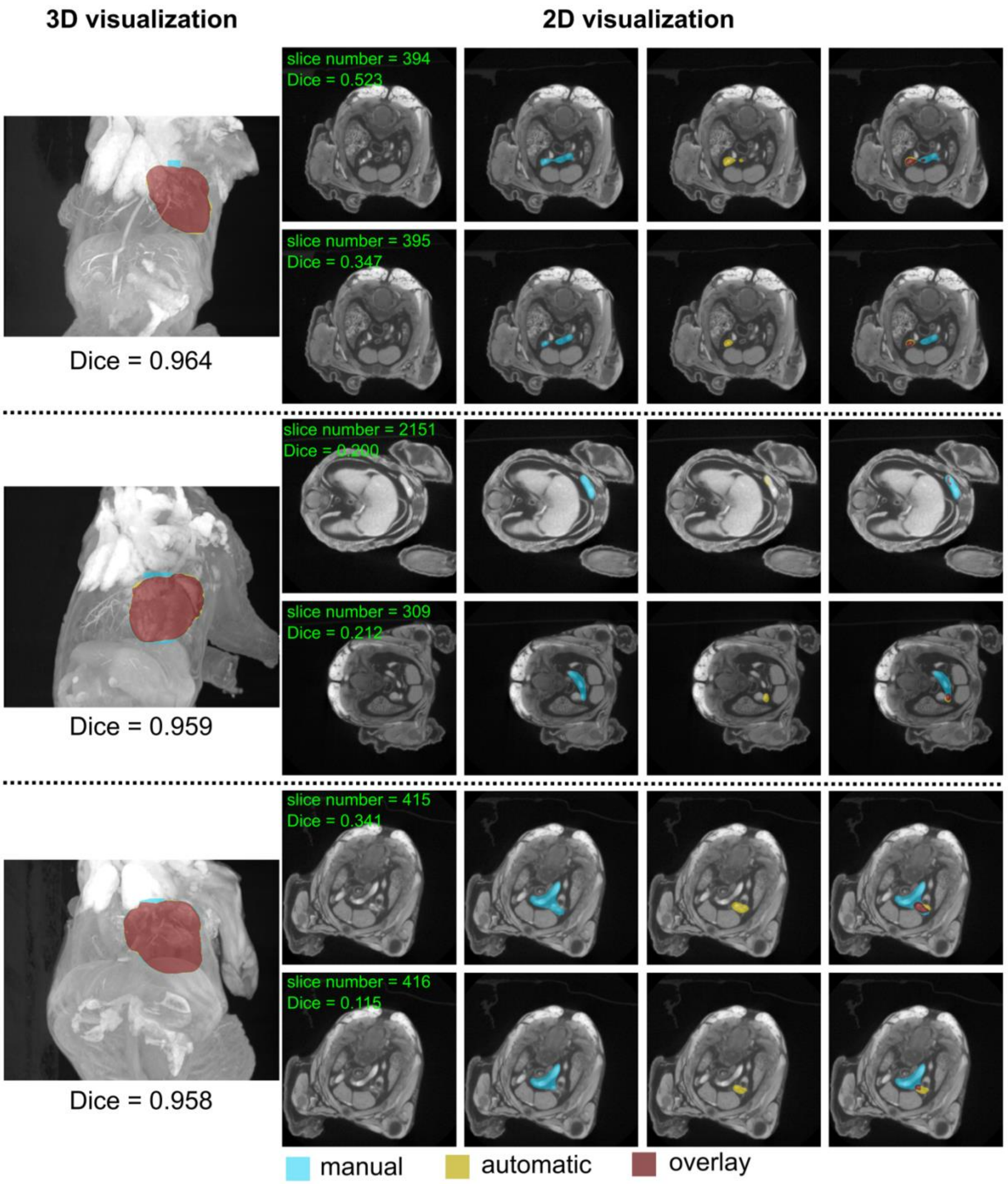

**Supplementary Figure 1: Examples of imperfectly segmented µCT scans.**

Three examples of incorrectly segmented µCT scans, (corresponding to the three lowest 3D Dice scores), based on comparison with a segmentation by an embryologist. 3D renderings are shown on the left, and two 2D slices with the lowest 2D Dice scores on the right. The four columns show the original image (grey), the manual segmentation (blue), the automated segmentation (yellow), and the overlap of manual and automated segmentations (red).

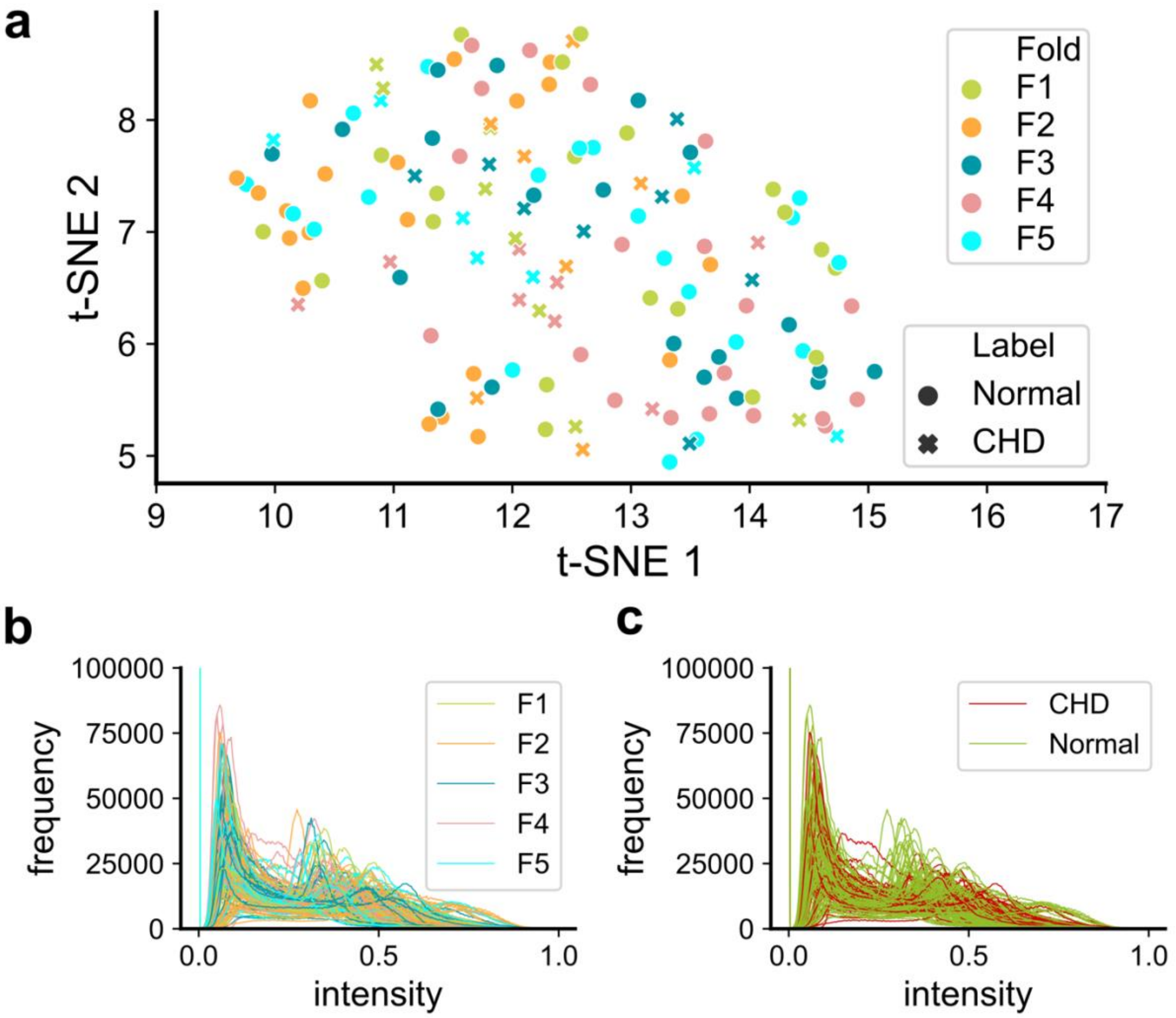

**Supplementary Figure 2: Distribution of scans and pixel intensities.**

**a)** t-distributed stochastic neighbor embedding (t-SNE) visualization of µCT images (each symbol corresponds to a distinct 3D image) for normal hearts (dots) and hearts with CHD (crosses). Colors indicate the data fold (F1-F5). **b**,**c)** Histograms of pixel intensities (each curve corresponds to a distinct 3D image), with one color for each data fold (F1-F5) (**b**) or one color for each classification (CHD vs normal) (**c**).

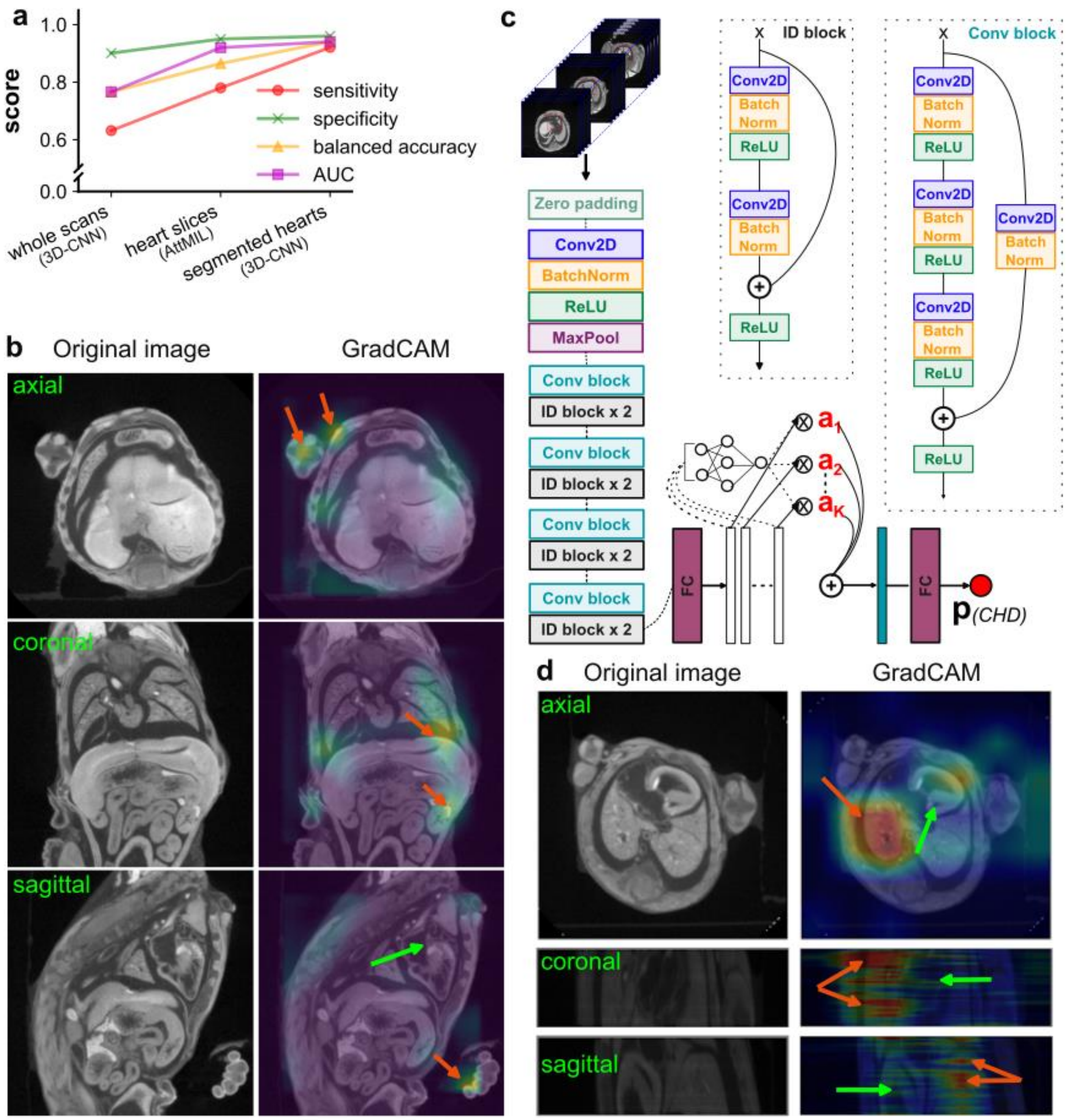

**Supplementary Figure 3: Diagnosis models without heart segmentation.**

**a)** Line chart compares the performance of three distinct classification models. From left to right: (i) the 3D-CNN model trained on entire body scans without heart segmentation ('whole scans'), (ii) an attention-based deep multiple instance learning (AttMIL) model applied to heart slices ('heart slices'), and (iii) the 3D-CNN model trained on segmented hearts ('segmented hearts'). **b)** For the segmentation-free 'whole scan' model, the GradCAM reveals important features (orange arrows) located outside the heart (green arrow). **c)** Architecture of AttMIL, utilizing ResNet50 as base model. The attention pooling module is introduced to measure the importance of each slice in the heart. The heart slices are resized to 224 x 224 pixels and fed into AttMIL to compute the probability of heart anomalies. **d)** For the AttMIL model, the GradCAM emphasizes the lungs (orange arrows) as important areas, rather than the heart (green arrows).

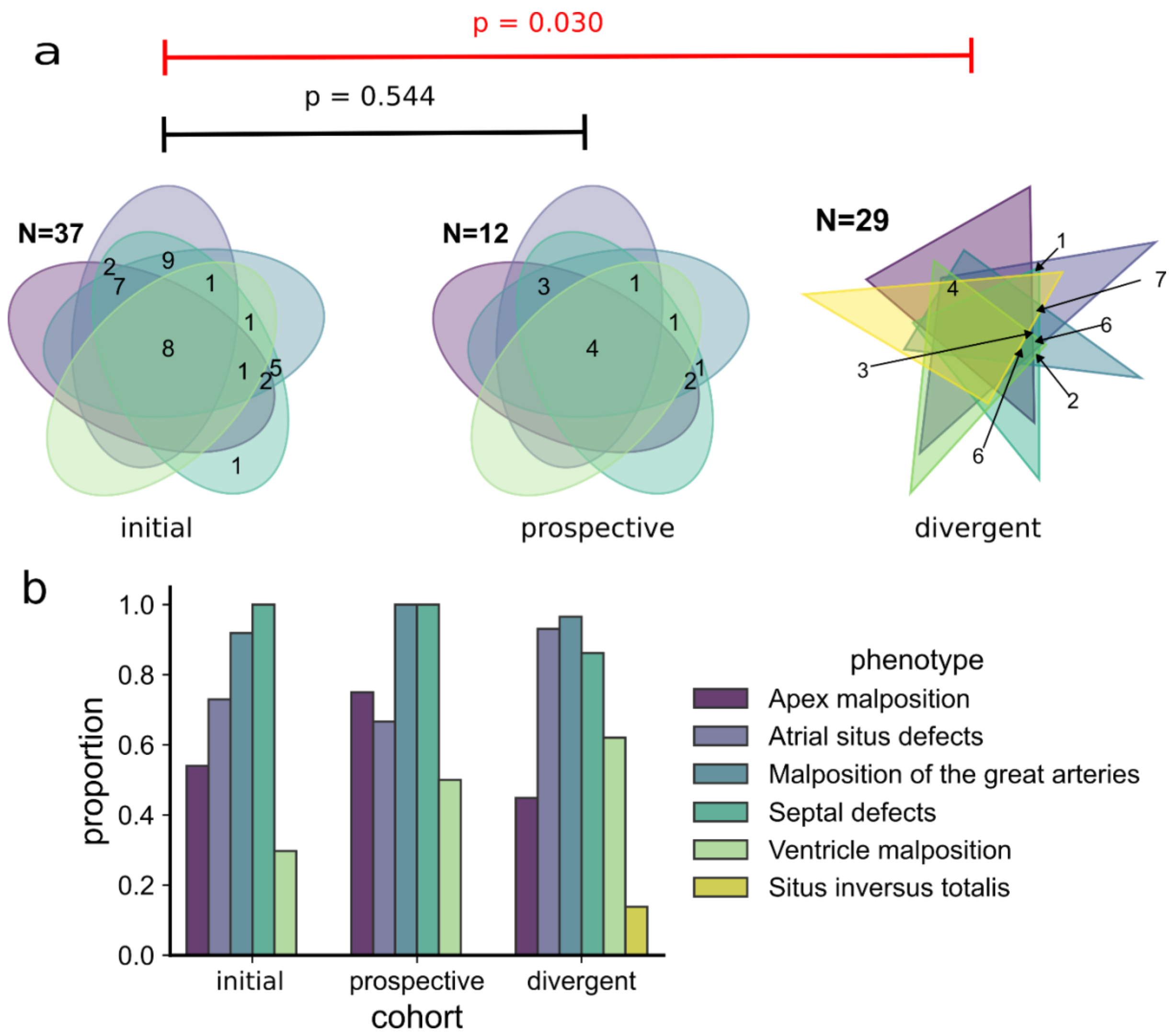

**Supplementary Figure 4: Phenotype distribution in different cohorts**

**a)** Venn diagrams show the combinations of phenotypes in the initial cohort (left), prospective cohort (middle), and the divergent cohort (right). A χ2 test shows no significant difference between combinations of phenotypes in the initial vs. prospective cohorts but shows a significant difference between the initial vs. divergent cohorts (p=0.03), with the presence of a new phenotype (situs inversus totalis, yellow). **b**) Proportions of the different phenotypes for each cohort.

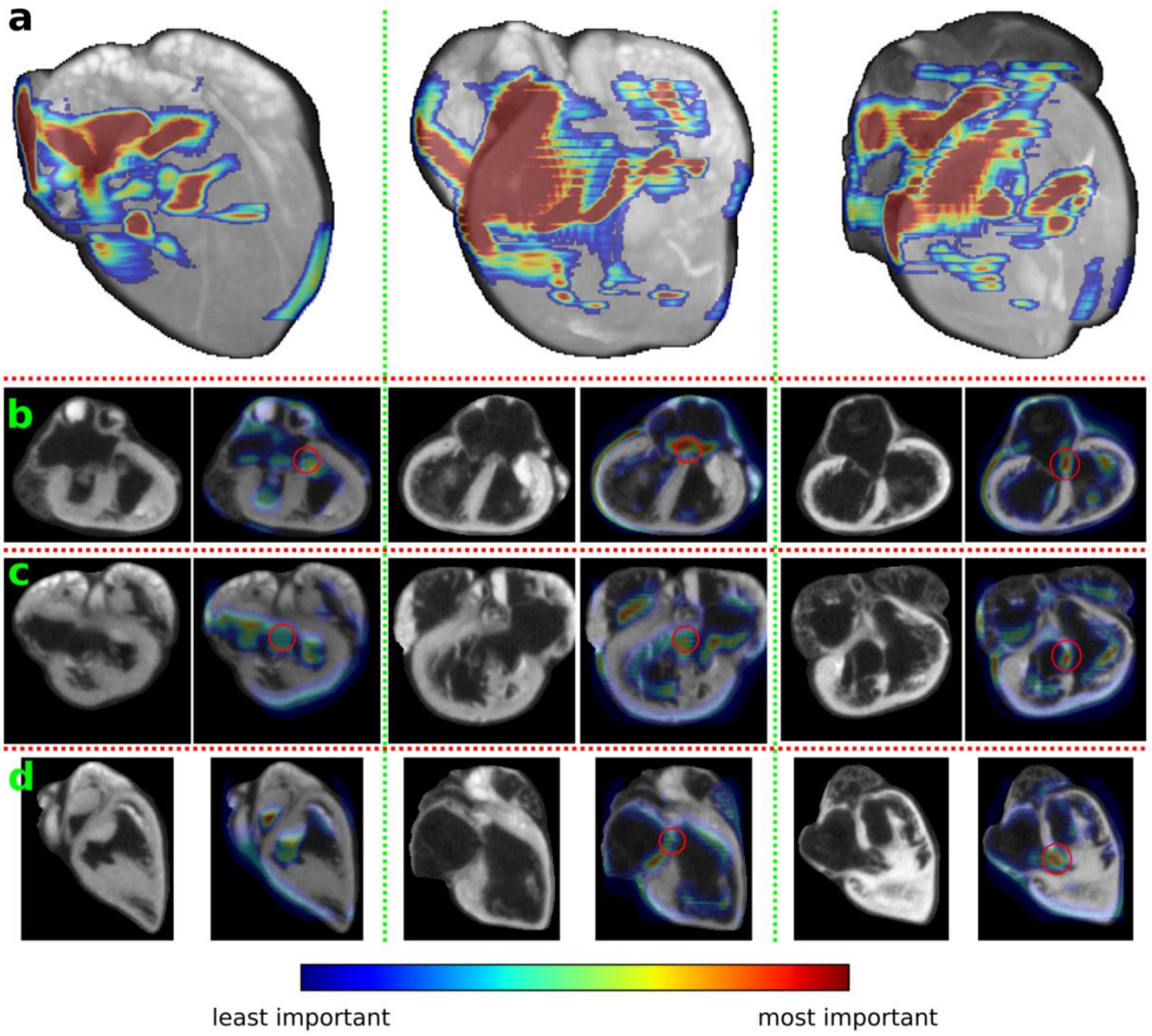

**Supplementary Figure 5: GradCAM visualizations of µCT scans with CHD**

Three scans were randomly chosen among those correctly predicted to have CHD. The Figure shows the original image (grey) with the GradCAM activation map for CHD superposed as a heat map. Hot colors indicate important regions for the classification output, cold colors indicate less important regions. **a**) 3D visualization. **b**) Axial view. **c**) Coronal view. **d**) Sagittal view. The atrioventricular junction is highlighted by the red circles. Out of 35 correctly predicted cases of CHD, 30 cases (86%) exhibited a GradCAM signal in the atrioventricular junction and septum (86%).

**Supplementary Table 1: Different mouse lines, genotypes, cardioplegia treatments and developmental stages across initial, prospective, and divergent cohorts.**

| | Initial cohort (n=139) | Prospective cohort (n=18) | Divergent cohort (n=80) |
|---|---|---|---|
| **Lines** | • *Nodal* conditional mutants (*Mef2cAHFCre*)<br>• *Nodal* conditional mutants (*Hoxb1*$^{Cre}$)<br>• *Nodal* ; *Ift20* double heterozygote mutants<br>• *Rpgrip1l* mutants | • *Nodal* conditional mutants (*Hoxb1*$^{Cre}$) | • *Ccdc40* mutants<br>• *Nodal* conditional mutants (*Hoxb1*$^{Cre}$) |
| **Genotypes** | • *Nodal*$^{flox/+}$<br>• *Nodal*$^{flox/+}$*; Tg Mef2cAHFCre*<br>• *Nodal*$^{flox/Nul}$<br>• *Nodal*$^{flox/+}$*;Hoxb1*$^{+/+}$<br>• *Nodal*$^{flox/Nul}$*;Hoxb1*$^{Cre/+}$<br>• *Nodal*$^{flox/+}$*;Hoxb1*$^{Cre/+}$<br>• *Ift20*$^{+/+}$*;Nodal*$^{+/+}$ (wild-types)<br>• *Ift20*$^{Nul/+}$*;Nodal*$^{+/+}$<br>• *Ift20*$^{Nul/+}$*;Nodal*$^{Nul/+}$<br>• *Ift20*$^{+/+}$*;Nodal*$^{Nul/+}$<br>• *Rpgrip1l*$^{+/+}$ (wild-types)<br>• *Rpgrip1l*$^{-/-}$ | • *Nodal*$^{flox/+}$*;Hoxb1*$^{+/+}$<br>• *Nodal*$^{flox/Nul}$*;Hoxb1*$^{Cre/+}$ | • *Ccdc40*$^{lnks/lnks}$<br>• *Ccdc40*$^{+/+}$ (wild-types)<br>• *Nodal*$^{flox/Nul}$*;Hoxb1*$^{Cre/+}$<br>• *Nodal*$^{flox/+}$*;Hoxb1*$^{+/+}$ |
| **Cardioplegia** | • KCl 250mM | • KCl 250mM | • 110mM NaCl, 16mM KCl,16mM $MgCl_2$, 1.5mM $CaCl_2$, 10mM $NaHCO_3$ |
| **Stages** | • E18.5<br>• P0 | • E18.5 | • E17.5<br>• E18.5<br>• P0 |

**Supplementary Table 2: Data partitioning of the prospective and divergent cohorts for retraining**

| | | Train | Test |
|---|---|---|---|
| **Mouse lines** | • *Ccdc40* mutants | 22 | 25 |
| | • *Nodal* conditional mutants (*Hoxb1*$^{Cre}$) *(**)* | 36 | 15 |
| **Genotypes** | • *Ccdc40*$^{lnks/lnks}$ | 12 | 15 |
| | • *Ccdc40*$^{+/+}$ (wild-types) | 10 | 10 |
| | • *Nodal*$^{flox/Nul}$*;Hoxb1*$^{Cre/+}$ *(**)* | 23 | 10 |
| | • *Nodal*$^{flox/+}$*;Hoxb1*$^{+/+}$ *(**)* | 13 | 5 |
| **Cardioplegia** | • *110mM NaCl, 16mM KCl,16mM* $MgCl_2$*, 1.5mM* $CaCl_2$*, 10mM* $NaHCO_3$ | 40 | 40 |
| | • *KCl 250mM (*)* | 18 | 0 |
| **Label** | • *Normal* | 31 | 26 |
| | • *CHD* | 27 | 14 |
| | **Total** | 58 | 40 |

*(*) only in prospective cohort*

*(**) both in prospective and divergent cohorts*
